# A Gene Co-expression Network-based Analysis of Multiple Brain Tissues Reveals Novel Genes and Molecular Pathways Underlying Major Depression

**DOI:** 10.1101/591693

**Authors:** Zachary F Gerring, Eric R Gamazon, Major Depressive Disorder Working Group of the Psychiatric Genomics Consortium, Eske M Derks

## Abstract

Major depression is a common and severe psychiatric disorder with a highly polygenic genetic architecture. Genome-wide association studies have successfully identified multiple independent genetic loci that harbour variants associated with major depression, but the exact causal genes and biological mechanisms are largely unknown. Tissue-specific network approaches may identify molecular mechanisms underlying major depression and provide a biological substrate for integrative analyses. We provide a framework for the identification of individual risk genes and gene co-expression networks using genome-wide association summary statistics and gene expression information across multiple human brain tissues and whole blood. We developed a novel gene-based method called eMAGMA that leverages multi-tissue eQTL information to identify 99 biologically plausible risk genes associated with major depression, of which 58 are novel. Among these novel associations is Complement Factor 4A (C4A), recently implicated in schizophrenia through its role in synaptic pruning during postnatal development. Major depression risk genes were enriched in gene co-expression modules in multiple brain tissues and the implicated gene modules contained genes involved in synaptic signalling, neuronal development, and cell transport pathways. Modules enriched with major depression signals were strongly preserved across brain tissues, but were weakly preserved in whole blood, highlighting the importance of using disease-relevant tissues in genetic studies of psychiatric traits. We identified tissue-specific genes and gene co-expression networks associated with major depression. Our novel analytical framework can be used to gain fundamental insights into the functioning of the nervous system in major depression and other brain-related traits.

**Author summary:** Although genome-wide association studies have identified genetic risk variants associated with major depression, our understanding of the mechanisms through which they influence disease susceptibility remain largely unknown. Genetic risk variants are highly enriched in non-coding regions of the genome and affect gene expression. Genes are known to interact and regulate the activity of one another and form highly organized (co-expression) networks. Here, we generate tissue-specific gene co-expression networks, each containing groups of functionally related genes or “modules”, to delineate interactions between genes and thereby facilitate the identification of gene processes in major depression. We developed and applied a novel research methodology (called “eMagma”) which integrates genetic and transcriptomic information in a tissue-specific analysis and tests for their enrichment in gene co-expression modules. Using this novel approach, we identified gene modules in multiple tissues that are both enriched with major depression genetic association signals and biologically meaningful pathways. We also show gene modules are strongly preserved across brain regions, but not in whole blood, suggesting blood may not be a useful tissue surrogate for the genetic dissection of major depression. Our novel analytical framework provides fundamental insights into the functional genetics major depression and can be applied to other neuropsychiatric disorders.

## Introduction

Major Depression is a highly disabling mental health disorder that accounts for a sizable proportion of the global burden of disease. The global lifetime prevalence of major depression is around 12% (17% of women and 9% of men) (1), and ranks as the fourth most disabling disorder in Australia in terms of years lived with disability (2). Major Depression has a complex molecular background, driven in part by a highly polygenic mode of inheritance. A recent genome-wide association study (GWAS) meta-analysis of 135,458 major depression cases and 344,901 controls identified 44 loci associated with the disorder (3). Detailed functional studies showed these loci to be enriched in multiple brain tissues and neuronal cell types and to contain common (minor allele frequency, MAF > 0.01) single nucleotide polymorphisms (SNPs) that regulate the expression of multiple genes with putative roles in central nervous system development and synaptic plasticity. These results suggest disease-associated SNPs modify major depression susceptibility by altering the expression of their target genes in the brain. Genes are known to interact and regulate the activity of one another in large gene-co-expression networks. Therefore, SNPs may not only affect the activity of single target gene, but multiple biologically related genes within the same tissue-specific co-expression network. The integration of GWAS SNP genotype data with gene co-expression networks across multiple tissues may be used to elucidate biological pathways and processes underlying highly polygenic complex disorders such as major depression.

Genome-wide gene expression data has been successfully integrated with SNP genotype data to prioritise risk genes and reveal possible mechanisms underlying susceptibility to a range of psychiatric disorders (4–6). This approach is most appropriate when performed in phenotypically affected cases and healthy controls for whom both gene expression and SNP genotype data are available. In practice, however, phenotype, SNP genotype, and gene expression data measured from the same individuals are difficult to obtain due to cost and tissue availability, and identifying causal variants can be difficult due to linkage disequilibrium (LD) and confounding from environmental and technical factors. Recent approaches address these limitations by integrating GWAS summary statistics with independent gene expression data provided by large international consortia, such as the multi-tissue Genotype-Tissue Expression (GTEx) project (7–9). The most recent release of the GTEx project (version 7) contains SNP genotype data linked to gene expression across 53 tissues from 714 donors, including 13 brain tissues from 216 donors. This represents a valuable resource with which to study gene expression and its relationship with genetic variation, known as expression quantitative trait loci (eQTL) mapping (10).

Recent genetic studies have leveraged GTEx data in gene-based analyses to prioritise individual risk genes whose expression is associated with major depression (11,12). While these analyses identified individual risk genes for major depression, they provide little insight into the molecular context within which the risk genes operate. We propose the use of GTEx data to build gene co-expression networks consisting of highly correlated modules—or groups—of genes in multiple tissues. The gene network modules provide a detailed map of gene interactions in a given tissue, and provide a biological substrate to test the enrichment of major depression GWAS signals. Enriched gene modules can be characterised using gene pathway analysis, and provide a valuable resource for the integration of additional molecular data. This approach may characterise the broader molecular context of risk genes in major depression and thereby facilitate the identification of gene pathways for diagnostic, prognostic, and therapeutic intervention.

## Results

### Genes form co-expression networks enriched in distinct biological processes

We built gene co-expression networks using RNA-Seq data from 13 brain tissues and whole blood in GTEx (v7). In total, 464 tissue samples (including 216 brain samples) and 17793 protein-coding genes were used to build the co-expression networks, although the number of samples (range: 80-369) and genes (range: 14834-16892) differed by tissue (Table 1). The number of gene co-expression modules within each gene network ranged from 11 modules in brain cortex to 24 modules in amygdala, and the number of genes within a module ranged from as few as 30 (0.18% of network genes, amygdala) to 9144 (55% of network genes, anterior cingulate cortex). We used gene pathway analysis to characterise biological processes in each co-expression module (Table S1). Co-expression networks were largely enriched for a single type of biological process (e.g. transcriptional regulation or immune response). These data suggest the network gene co-expression modules represent biologically homogeneous units.

**Table 1:** Summary of GTEx gene expression information used to build the gene-expression networks with descriptive statistics and P value thresholds for gene-based analyses

| Tissue | Gene Network |  | Modules | Gene Module Size |  |  | eMAGMA |  | S-PrediXcan |  |
| --- | --- | --- | --- | --- | --- | --- | --- | --- | --- | --- |
|  | Samples | Genes | N | Min | Median | Max | Genes | Threshold | Genes | Threshold |
| Amygdala | 88 | 16547 | 24 | 30 | 362 | 4125 | 1214 | 4.12E-05 | 2338 | 2.14E-05 |
| Anterior cingulate cortex | 109 | 16568 | 16 | 55 | 391 | 9144 | 2356 | 2.12E-05 | 3269 | 1.53E-05 |
| Caudate basal ganglia | 144 | 16612 | 16 | 75 | 483 | 6971 | 3272 | 1.53E-05 | 4142 | 1.21E-05 |
| Cerebellar Hemisphere | 125 | 16505 | 13 | 163 | 741 | 8662 | 4075 | 1.23E-05 | 4726 | 1.06E-05 |
| Cerebellum | 154 | 16607 | 15 | 86 | 613 | 8133 | 5386 | 9.28E-06 | 6044 | 8.27E-06 |
| Cortex | 136 | 16665 | 11 | 126 | 582 | 6188 | 3625 | 1.38E-05 | 4299 | 1.16E-05 |
| Frontal Cortex | 118 | 16642 | 22 | 51 | 345 | 4965 | 2952 | 1.69E-05 | 3565 | 1.40E-05 |
| Hippocampus | 111 | 16634 | 17 | 84 | 388 | 4168 | 1830 | 2.73E-05 | 2788 | 1.79E-05 |
| Hypothalamus | 108 | 16892 | 17 | 60 | 613 | 5322 | 1605 | 3.12E-05 | 2819 | 1.77E-05 |
| Nucleus accumbens | 130 | 16652 | 20 | 68 | 471 | 5346 | 2842 | 1.76E-05 | 3593 | 1.39E-05 |
| Putamen | 111 | 16399 | 15 | 53 | 695 | 7371 | 2401 | 2.08E-05 | 3153 | 1.59E-05 |
| Spinal cord cervical c-1 | 83 | 16809 | 20 | 56 | 577 | 3017 | 1470 | 3.40E-05 | 2501 | 2.00E-05 |
| Substantia nigra | 80 | 16612 | 19 | 73 | 475 | 5266 | 989 | 5.06E-05 | 2015 | 2.48E-05 |
| Whole Blood | 369 | 14834 | 14 | 57 | 293 | 6939 | 5946 | 8.41E-06 | 6249 | 8.00E-06 |

### Identification of risk genes for major depression

To assign major depression risk SNPs to genes, we applied two gene-based strategies: first, proximity-based gene mapping with MAGMA, which assigns SNPs to the nearest gene within a genomic window; and second, eQTL gene mapping using eMAGMA, which uses tissue-specific SNP-gene associations from GTEx to assign SNPs to genes based on their association with gene expression. To further prioritise gene-level results, we performed a transcriptome-wide association study using S-PrediXcan. Both tissue-specific and *P* value thresholds for each gene-based method, calculated using Bonferroni correction for the number of associations, are shown in Table 1. We identified 137 unique mapped depression-associated genes with MAGMA (Table S2), 217 significant tissue-specific gene associations with eMAGMA (representing 99 unique mapped genes) (Table S3), and 86 tissue-specific gene associations with S-PrediXcan (Table S4). A total of 41 genes were implicated by both MAGMA and eMAGMA mapping strategies in at least one tissue (Figure 1; Table S5). Among significant eMAGMA associations, 35 (16%) also had a significant S-PrediXcan association in the same tissue (Table S6), and 16 associations were significant across all three gene-based methods (Table 2). Taken together, these results point to potential functional links for the GWAS-associated variants and give higher credibility to genes with convergent evidence of association from multiple sources.

**Figure 1:**
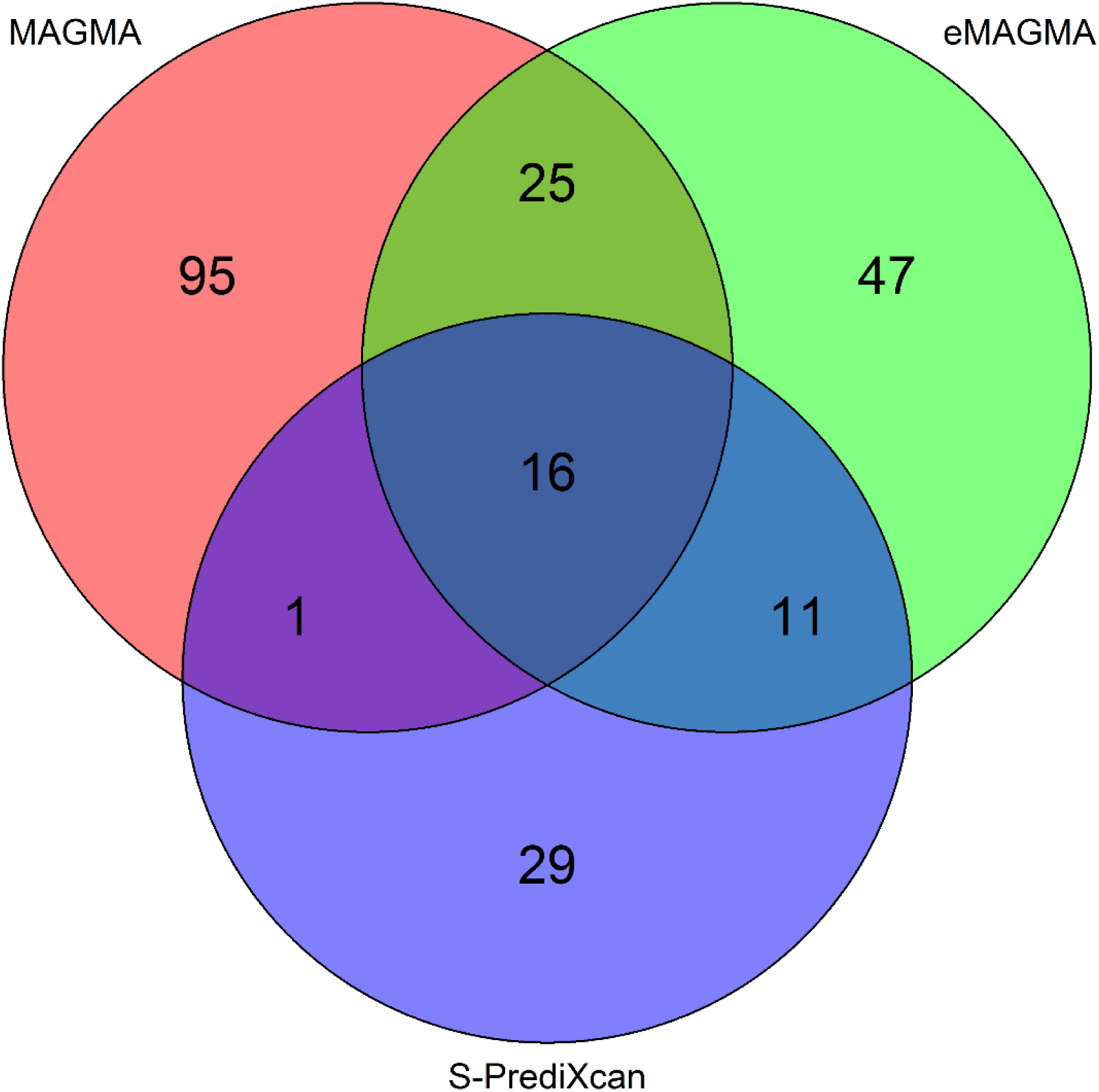
Overlap in the number of significant gene-level associations between MAGMA, eMAGMA, and S-PrediXcan. **Notes:** Significant MAGMA genes (N=137) were selected using Bonferroni correction for the entire list of gene-based *P* values (i.e. 0.05/18042=2.77×10^−6^). Significant eMAGMA (N=99) and S-PrediXcan (N=57) results were adjusted using Bonferroni correction for the number of associations in each tissue (see Table 1 for tissue-specific thresholds). The overlap between gene-level results after correcting for all tissues and genes (N=51,501, *P*=9.71×10^−7^) is presented in Figure S1. Refer to Table 3 for the top (N=10) overlapping significant gene-based associations for MAGMA, eMAGMA, and S-PrediXcan, and Table S5 for the entire list of gene-based results (N=41).

**Table 2:**
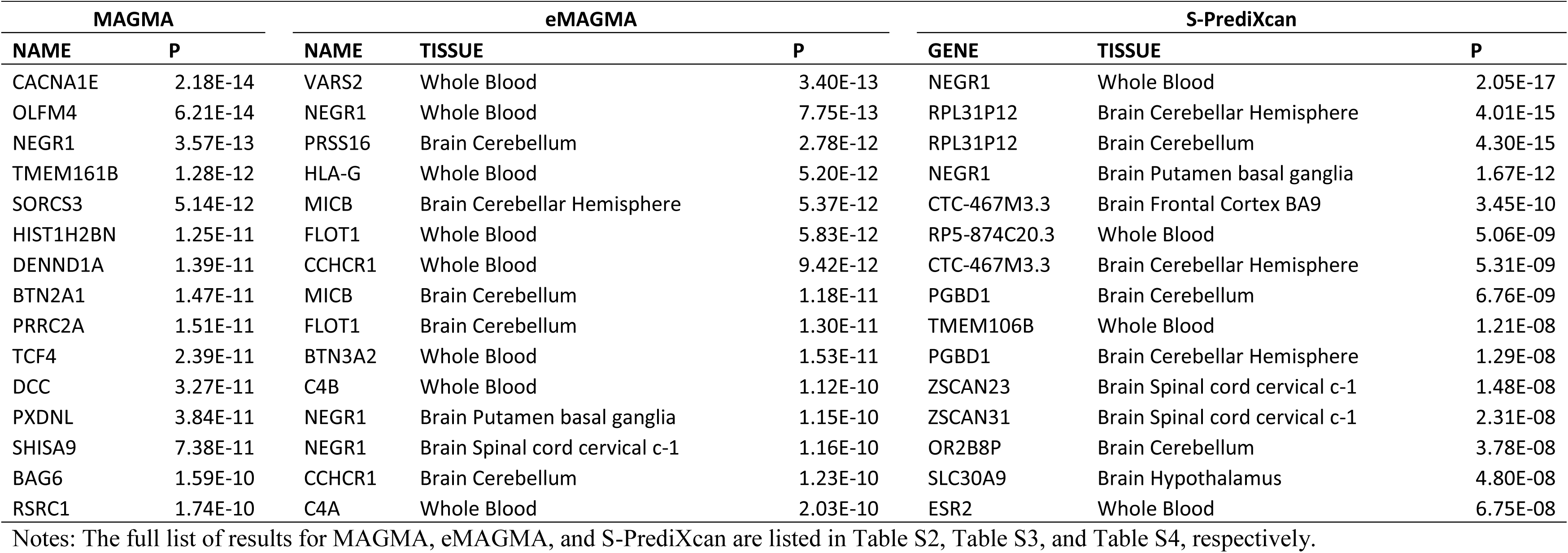
Significant major depression candidate risk genes for three gene-based methods (MAGMA, eMAGMA, S-PrediXcan)

**Table 3:** Major Depression association signals are enriched in gene co-expression network modules.

| Module | Tissue | Genes (N) | Beta | SE | P | P corr |
| --- | --- | --- | --- | --- | --- | --- |
| MAGMA |  |  |  |  |  |  |
| M1 | Amygdala | 3681 | 0.0599 | 0.0172 | 0.0002 | 0.0055 |
| M2 | Frontal Cortex | 4507 | 0.0532 | 0.0162 | 0.0005 | 0.0127 |
| M3 | Nucleus accumbens | 4817 | 0.0468 | 0.0158 | 0.0015 | 0.0285 |
| M4 | Cerebellar Hemisphere | 994 | 0.0822 | 0.0304 | 0.0035 | 0.0433 |
| eMAGMA |  |  |  |  |  |  |
| M5 | Hypothalamus | 20 | 0.7480 | 0.2510 | 0.0015 | 0.0267 |
| M6 | Putamen | 1062 | 0.1230 | 0.0434 | 0.0023 | 0.0392 |

**Notes:** Module enrichment analyses was performed using MAGMA. Beta: regression coefficient of the gene set; SE, the standard error of the regression coefficient; P the competitive gene-set P value; P corr: P value empirically corrected for multiple testing for all the gene set.

### Major depression risk genes are enriched in brain gene co-expression network modules

We tested for the enrichment of MAGMA (Table S2; N=137) and eMAGMA associations (Table S3; N unique=99) in gene co-expression modules from the brain and whole blood. Gene modules in four brain tissues (amygdala, cerebellar hemisphere, frontal cortex, and nucleus accumbens) were enriched with MAGMA association signals, while one module in hypothalamus and one module in putamen were enriched with eMAGMA associations (Table 4). Gene modules enriched with MAGMA remained significant after removal of genes in the MHC region (Table S7), however modules enriched with eMAGMA associations were no longer significant after empirical multiple testing correction (Table S8). No enrichment of gene-based association signals was observed for modules identified in whole blood, despite the larger sample size (and hence increase power) compared to brain tissues. We plotted the overlap in gene modules enriched with gene-based MD associations (Figure S1). A total of 217 genes overlapped across four modules enriched with MAGMA associations, suggesting similar biological processes may underlie the modular enrichments. On the other hand, we observed little overlap in genes between modules enriched eMAGMA associations, suggesting tissue-specific eQTL effects may underlie these modular enrichments and highlighting the importance of using both proximity-and eQTL-based gene-based tests.

In gene pathway analyses of the major depression enriched modules, we found enrichment of neuronal and synaptic signalling pathways in amygdala, frontal cortex, nucleus accumbens, putamen (e.g. trans-synaptic signalling in frontal cortex, P=2.81 × 10^−24^), as well as membrane trafficking related pathways in cerebellar hemisphere (e.g. Membrane Trafficking, P=2.19 × 10^−13^) and vascular-related pathways in hypothalamus (e.g. blood vessel morphogenesis, P=5.67 × 10^−15^) (Figure 2, Table S9). Pathway analysis of 217 genes overlapping four modules enriched with MAGMA gene-based associations revealed chemical synaptic transmission (GO:0007268; *P*=1.24 × 10^−14^) and the neuronal system (R-HSA-112316; *P*=6.62 × 10^−10^) pathways (Table S10).

**Figure 2:**
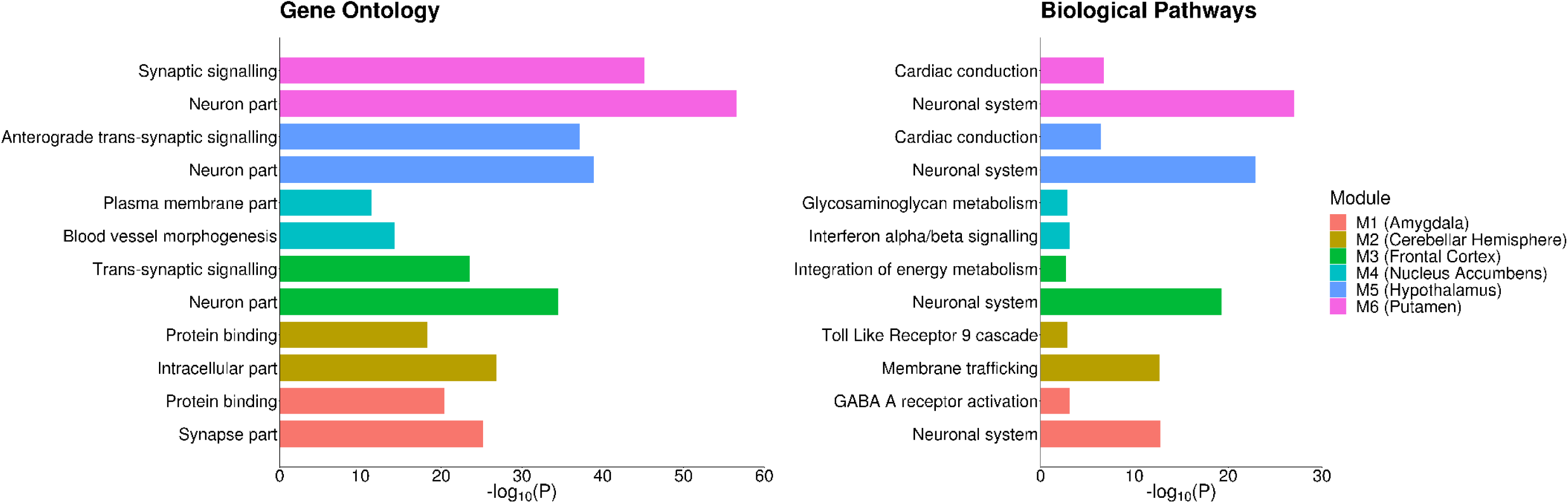
Pathway analysis of major depression enriched modules **Notes:** A competitive gene pathway analysis was performed on tissue-specific significant gene co-expression modules using g:Profiler (https://biit.cs.ut.ee/gprofiler/index.cgi). The figure shows the gene ontology and biological pathways of tissue-specific modules enriched with major depression gene-based signals.

**Figure 3:**
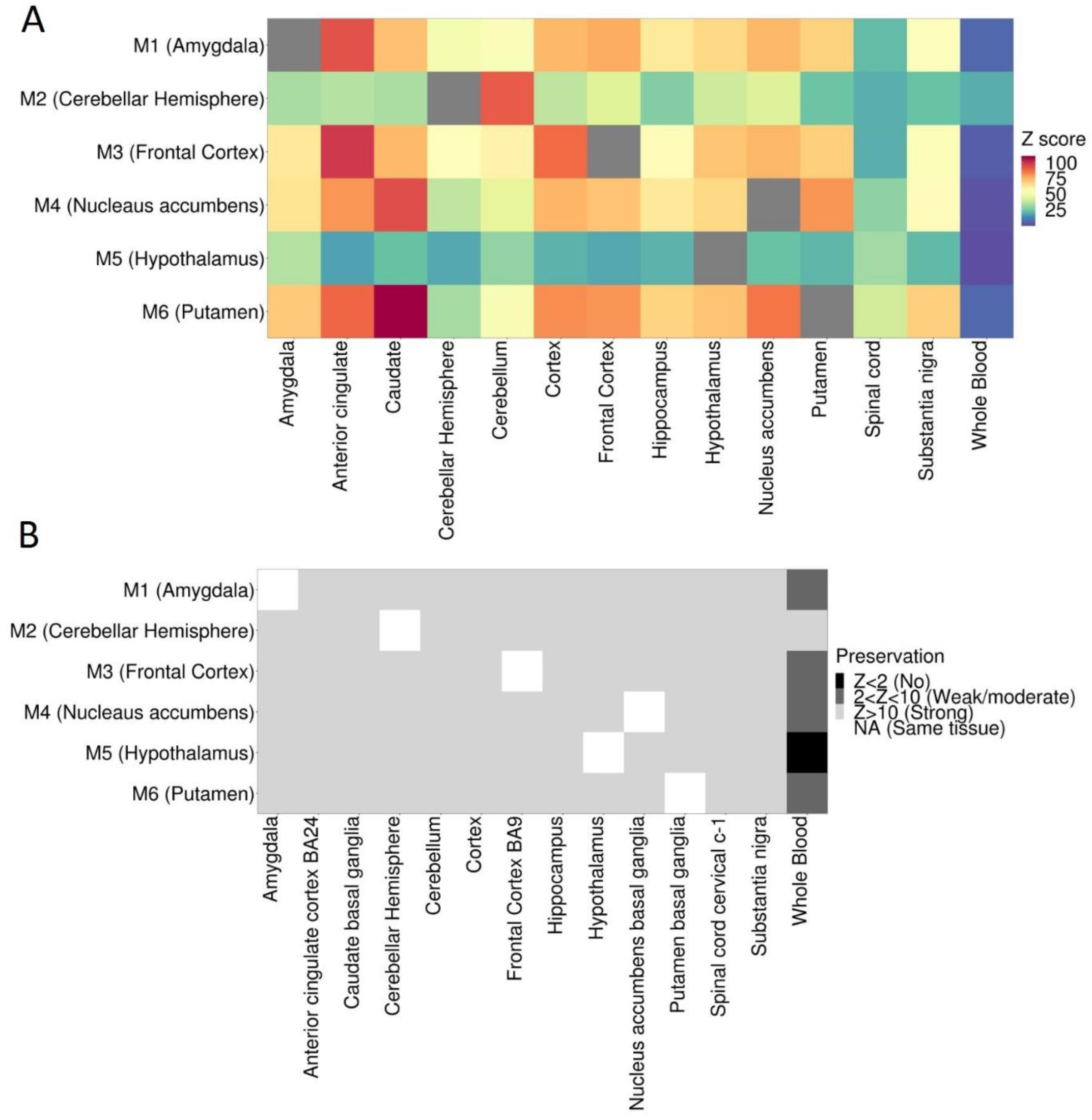
Preservation of major depression enriched network modules across brain tissues and whole blood. **(A)** Preservation Z score for tissue-specific modules (labelled M1 to M6 on the y axis) across brain tissues and whole blood. A higher Z score indicates greater preservation (i.e. replication) of a “reference” network in a “test” network (and vice versa). **(B)** Categorical classification of preservation Z scores across brain tissues and whole blood. A Z score less than 2 indicates no modular preservation; a Z score between 2 and 10 indicates weak to moderate preservation; and a Z score greater than 10 indicates strong preservation.

### Gene co-expression modules enriched with major depression risk genes are preserved across brain tissues

Our network-based approach allows the discovery of major depression associated gene modules as well as the preservation (or reproducibility) of those associated modules across tissues. We assessed the preservation of gene co-expression modules across brain tissues and whole blood using the WGCNA *modulePreservation* algorithm, highlighting the preservation of modules enriched with major depression GWAS signals. Strong modular preservation (Z score > 10) was observed across all brain regions, while weak to moderate preservation was observed in whole blood (Z score < 10) (Supplementary Figure 2). Major depression modules enriched with synaptic signalling pathways (Modules M1 [Amygdala], M3 [Frontal cortex], M4 [Nucleus accumbens], and M6 [Putamen]) showed particularly strong preservation across brain tissues, while module M2 (cerebellar hemisphere), enriched with cellular localisation and transport pathways, and module M5 (Hypothalamus), enriched with vascular related pathways, showed relatively weak preservation (Figure 2).

## Discussion

Genome-wide association studies have provided important insight into the genetic architecture of major depression. The next critical step is to leverage these genetic data to identify higher order biological processes underlying the disorder, and to ultimately identify molecular targets for risk prediction, diagnosis, and therapeutic intervention. To this end, we integrated multi-tissue gene expression data with major depression GWAS summary statistics using an integrative network-based approach. We first applied a weighted gene co-expression network analysis to gene expression data from multiple tissues in GTEx to measure the correlation structure between protein-coding genes. The gene networks, which represent the “connectedness” of genes in a given tissue, were divided into modules (or groups) of highly correlated genes, under the assumption that correlated genes are involved in the same or similar biological processes. The gene modules subsequently formed the unit of analysis for a) tests of enrichment with major depression GWAS summary statistics; b) gene pathway analyses using curated gene sets; and c) modular preservation (or replication) analyses across multiple tissues. Our network-based approach identified novel gene candidates for major depression, and identified co-expression networks enriched with biologically meaningful gene pathways that were preserved across brain tissues.

A gene module approach based on gene co-expression patterns was used to identify groups of functionally related genes associated with major depression. This approach reduced the dimensionality of genome-wide gene expression data across multiple brain tissues and whole blood without the loss of important biological information, and thereby alleviated the multiple testing burden associated with traditional single gene-based methods. A similar network-based approach has been applied to gene expression data for other brain-related disorders, including post-traumatic stress syndrome (13), schizophrenia (14), and psychosis (15). However, these studies typically included a small number of individuals (fewer than 100) from a single brain region and are therefore limited in their statistical power and generalisability across different tissues. Our approach used a total of 216 individuals with a tissue sample from at least one of 13 brain regions, and 464 individuals with the inclusion of whole blood, thereby improving the resolution and robustness of gene networks. Our network approach identified between 11 (Cortex) and 24 (Amygdala) mutually exclusive modules within tissues, and ranged in size from 30 to 9144 genes. Each module was enriched with distinct and highly significant biological pathways (e.g. immune signalling), suggesting our approach generated robust modules of functionally related genes.

To identify genes and gene-sets associated with major depression, we assigned disease-associated SNPs to their nearest gene using both proximity and multi-tissue eQTL information. We first used MAGMA, a proximity-based approach that assigns SNPs to their nearest gene. This approach appropriately corrects for correlated SNPs (i.e. linkage disequilibrium [LD]), and also adjusts for correlated gene expression in gene-set analysis and multiple-testing correction. However, SNPs are simply assigned to their nearest gene based on an arbitrary genomic window. It is well known that such proximity-based approaches often miss the functional SNP-gene association (16). Therefore, we modified the MAGMA pipeline to integrate SNPs with a significant (FDR<0.05) association with the expression of one or more genes (eQTLs) from the multi-tissue GTEx resource. In doing so, we generated a tissue-specific, eQTL-informed MAGMA, or “eMAGMA”, for the functional (i.e. biologically meaningful) annotation of GWAS summary statistics.

Our eMAGMA approach identified novel and biologically meaningful candidate risk gene associations for major depression across multiple tissues. Of 99 significant eMAGMA genes (representing 217 unique gene-tissue associations), 58 were not identified by (proximity-based) MAGMA. Noteworthy among these associations is Complement Factor 4A (*C4A*), recently implicated in the development of schizophrenia through its role as a mediator of synaptic pruning during postnatal development (17). *C4A* was significant in 12 of 14 investigated tissues, including whole blood, and was one of 24 significant eMAGMA genes located on chromosome 6p21—a region with complex LD structure that flanks the centromeric end of the major histocompatibility complex. While we cannot identify the causal gene due to the complex LD pattern of the MHC region, this result suggests partially shared biological pathways underlie both major depression and schizophrenia.

We tested for the enrichment of candidate risk genes in tissue-specific network co-expression modules, while adjusting for correlated gene expression, gene size, and gene density. We identified six co-expression modules across six individual brain tissues, four of which were enriched in synaptic signalling and neuronal development pathways (Amygdala, Putamen, Frontal cortex, and hypothalamus). These results align with recent pathway analyses of genetic associations in major depression, which identified genes and gene-sets involved in synaptic transmission and neuronal mechanisms, among other pathway groupings (3,18). Furthermore, structural changes in frontal cortex have been identified in a recent meta-analysis of brain magnetic resonance imagining findings in adult major depression cases (19), highlighting the central role of frontal cortex in major depression aetiology. It is important to note that co-expression networks across all (N=13) brain tissues contained a gene co-expression module enriched with synaptic and neuronal pathways, but only four were enriched with major depression association risk genes. This suggests risk genes underlying major depression susceptibility manifest their effect in specific brain regions, consistent with tissue-specific gene expression (20) and highlighting the importance of studying multiple tissues in integrated studies of complex traits such as major depression.

We assessed the preservation (or reproducibility) of connectivity patterns (i.e. correlations) between genes across multiple brain tissues and whole blood. This approach may determine whether the connectivity between genes within a network module enriched with major depression signals differs both across brain tissues and between brain and whole blood, and may therefore identify (peripheral) surrogate tissues for molecular studies of major depression. We observed strong preservation of network modules across all brain regions, but not whole blood, suggesting blood-based molecular studies of major depression may fail to capture important disease-related processes in brain. Our findings therefore support the use of brain tissues from large international consortia, such as the GTEx study or the CommonMind consortium, for the characterisation of genetic association signals, despite reduced sample sizes compared to blood-based datasets and the potential for technical biases associated with the use of post-mortem samples.

Current results support a common variant genetic architecture of major depression, where variants with relatively high frequency (e.g. minor allele frequency > 0.01) in the general population, but low penetrance, are the major contributors to genetic susceptibility to the disorder. Therefore, as sample sizes grow larger, thousands of lead SNPs associated with major depression are likely to be identified. With these impending data, new methods for the interpretation of genetic signals for major depression and other common complex disorders will be required. Our network-based approach provides a gene expression substrate across multiple human tissues for the integration and characterisation of GWAS signals. By exploiting the connectivity between genes, this approach will allow the identification of perturbations in the activity of a system rather than individual genes. Furthermore, network-based methods may identify regulatory hubs whose perturbation may have wider consequences for major depression and other (co-morbid) psychiatric and/or neurological disorders by virtue of their interaction with other genes. Finally, gene co-expression networks can be integrated with other molecular phenotypes, such as epigenetic DNA methylation, proteomics, and metabolomics data, for ‘omics-based analyses. Such ‘omics-based analyses are critical for understanding the properties of biological systems underlying complex disorders such as major depression.

## Methods

### The Genotype-Tissue Expression (GTEx) project

An overview of our analytical pipeline is shown in supplementary Figure 1. Fully processed, filtered and normalised gene expression data for 13 brain tissues and whole blood (Table 1) were downloaded from the Genotype-Tissue Expression project portal (version 7) (http://www.gtexportal.org) (Table 1). Only genes with ten or more donors with expression estimates > 0.1 Reads Per Kilobase of transcript (RPKM) and an aligned read count of six or more within each tissue were considered significantly expressed. Within each tissue, the distribution of RPKMs in each sample was quantile-transformed using the average empirical distribution observed across all samples. Expression measurements for each gene in each tissue were subsequently transformed to the quantiles of the standard normal distribution.

### Genome-wide association study of major depressive disorder

Detailed methods, including a description of population cohorts, quality control of raw SNP genotype data, and association analyses for the major depression GWAS is described elsewhere (11). The major depression GWAS included a mega-analysis of 29 samples (PGC29) (16,823 major depression cases and 25,632 controls) of European ancestry and additional analyses of six independent European ancestry cohorts (118,635 cases and 319,269 controls). Cases in the PGC29 cohort satisfied diagnostic criteria (DSM-IV, ICD-9, or ICD-10) for lifetime major depression. Cases in the expanded cohort were collated using a variety of methods: Generation Scotland employed direct interviews; iPSYCH (Denmark) used national treatment registers; deCODE (Iceland) used national treatment registers and direct interviews; GERA used Kaiser-Permanente (health insurance) treatment records (CA, US); UK Biobank combined self-reported major depression symptoms and/or treatment for major depression by a medical professional; and 23andMe used self-report of treatment for major depression by a medical professional. Controls in PGC29 were screened for the absence of major depression. A combination of polygenic scoring and linkage disequilibrium score regression showed strong genetic homogeneity between the PGC29 and additional cohorts and between samples within each cohort. SNPs and insertion-deletion polymorphisms were imputed using the 1000 Genomes Project multi-ancestry reference panel (21). Logistic regression association tests were conducted for imputed marker dosages with principal components covariates to control for population stratification. Ancestry was evaluated using principal components analysis applied to directly genotyped SNPs. Summary statistics for 10,468,942 autosomal SNPs were made available by the PGC and were utilized in our study.

### Identification of gene expression modules

Gene co-expression modules were individually constructed for 13 brain tissues and whole blood using the weighted gene co-expression network analysis (WGCNA) package in R (22). An unsigned pairwise correlation matrix – using Pearson’s product moment correlation coefficient – was calculated. An appropriate “soft-thresholding” value, which emphasises strong gene-gene correlations at the expense of weak correlations, was selected for each tissue by plotting the strength of correlation against a series (range 2 to 20) of soft threshold powers. The correlation matrix was subsequently transformed into an adjacency matrix, where nodes correspond to genes and edges to the connection strength between genes. Each adjacency matrix was normalised using a topological overlap function. Hierarchical clustering was performed using average linkage, with one minus the topological overlap matrix as the distance measure. The hierarchical cluster tree was cut into gene modules using the dynamic tree cut algorithm (23), with a minimum module size of 30 genes. We amalgamated modules if the correlation between their eigengenes – defined as the first principal component of their genes’ expression values – was greater or equal to 0.8.

### Gene-level analysis of Major Depression GWAS signals

We identified and prioritised risk genes for major depression using three approaches. First, we performed gene-level analyses using MAGMA v1.06 (24). This approach assigns SNPs to their nearest gene using a pre-defined genomic window (here a 35 kb upstream or 10 kb downstream of a gene body) and computes a gene-based statistic based on the sum of the assigned SNP – log(10) *P* values while accounting for the correlation (i.e. linkage disequilibrium) between nearby SNPs. Second, we modified the MAGMA approach by integrating eQTL information from the GTEx project. That is, for a given interrogated tissue, we assigned SNPs to target genes based on significant (FDR<0.05) SNP-gene associations in GTEx. This approach, which we will refer to as “eMAGMA”, is a tissue-specific, eQTL-informed method for assigning SNPs to genes. Gene-based statistics were subsequently computed using the sum of the assigned SNP –log(10) *P* values, in a similar manner to proximity-based MAGMA. Third, we used S-PrediXcan to integrate eQTL information from GTEx with major depression GWAS summary statistics to identify genes whose genetically predicted expression levels are associated with major depression. For S-PrediXcan, we used expression weights for 13 brain tissues and whole blood generated from GTEx (v7) (25), and LD information from the 1000 Genomes Project Phase 3 (26). These data were processed with beta values and standard errors from the GWAS of major depression (3) to estimate the expression-GWAS association statistic. For each gene-level approach, we corrected for multiple testing using Bonferroni correction. For MAGMA, we corrected for the total number of genes tested (i.e. 0.05/18,041 = 2.77 × 10^−6^). For the multi-tissue eMAGMA and S-PrediXcan, we applied two correction thresholds (Table 1): a “liberal” threshold, which corrected for the number of tests within each tissue (i.e. ignoring the number of tissues tested), and a “conservative” threshold, which corrected for the total number of tests performed (i.e. all tests across all tissues).

### Gene-set analysis of gene co-expression modules

To identify gene co-expression modules enriched with major depression risk genes, we performed gene-set analysis of both (proximity) MAGMA and eMAGMA gene-level results in tissue-specific gene co-expression modules using the gene-sets analysis function in MAGMA v1.06. The competitive analysis tests whether the genes in a gene-set (i.e. gene co-expression module) are more enriched with major depression risk genes than other genes while accounting for gene size and gene density. We applied an adaptive permutation procedure (24) (N=10,000 permutations) to obtain P values corrected for multiple testing. The 1000 Genomes European reference panel (Phase 3) was used to account for Linkage Disequilibrium (LD) between SNPs. For each tissue and gene-based enrichment method, a quantile-quantile plot of observed versus expected P values was generated to assess inflation in the test statistic. Gene-set enrichment analyses were re-performed after excluding genes in the MHC region.

### Characterisation of gene expression modules

Gene expression modules enriched with major depression GWAS association signals were assessed for biological pathways and processes using g:Profiler (https://biit.cs.ut.ee/gprofiler/) (27). Ensembl gene identifiers within enriched gene modules were used as input; we tested for the over-representation of module genes in Gene Ontology (GO) biological processes, as well as KEGG(28) and Reactome(29) gene pathways. The g:Profiler algorithm uses a Fisher’s one-tailed test for gene pathway enrichment; the smaller the P value, the lower the probability a gene belongs to both a co-expression module and a biological term or pathway purely by chance. Multiple testing correction was done using g:SCS; this approach accounts for the correlated structure of GO terms and biological pathways, and corresponds for an experiment-wide threshold of α=0.05.

### Preservation of gene co-expression networks across tissues

To examine the tissue-specificity of biological pathways, we assessed the preservation (i.e. replication) of network modules across GTEx tissues using the “modulePreservation” R function implemented in WGCNA (30). Briefly, the module preservation approach takes as input “reference” and “test” network modules and calculates statistics for three preservation classes: i) density-based statistics, which assess the similarity of gene-gene connectivity patterns between a reference network module and a test network module; ii) separability-based statistics, which examine whether test network modules remain distinct in reference network modules; and iii) connectivity-based statistics, which are based on the similarity of connectivity patterns between genes in the reference and test networks. For simplicity, we report two density and connectivity composite statistics: “Zsummary” and “medianRank”. A Zsummary value greater than 10 suggests there is strong evidence a module is preserved between the reference and test network modules, while a value between 2 and 10 indicates weak to moderate preservation and a value less than 2 indicates no preservation. The median rank statistic ranks the observed preservation statistics; modules with lower median rank tend to exhibit strong preservation than modules with higher median rank.

## Acknowledgements

We are deeply indebted to the investigators who comprise the Psychiatric Genomics Consortium (PGC), and to the hundreds of thousands of subjects who have provided data. Collaborators for the 23andMe Research Team are: Michelle Agee, Babak Alipanahi, Adam Auton, Robert K. Bell, Katarzyna Bryc, Sarah L. Elson, Pierre Fontanillas, Nicholas A. Furlotte, David A. Hinds, Karen E. Huber, Aaron Kleinman, Nadia K. Litterman, Matthew H. McIntyre, Joanna L. Mountain, Elizabeth S. Noblin, Carrie A.M. Northover, Steven J. Pitts, J. Fah Sathirapongsasuti, Olga V. Sazonova, Janie F. Shelton, Suyash Shringarpure, Chao Tian, Joyce Y. Tung, Vladimir Vacic, Catherine H. Wilson.

## Supporting information captions

**Supplementary material 1:** Major Depressive Disorder Working Group of the Psychiatric Genomics Consortium

**Table S1:** Pathway enrichments of tissue-specific network modules

**Table S2:** Significant (Bonferroni correction, *P*<2.6 × 10^−6^) MAGMA gene-based results

**Table S3:** Significant (Bonferroni correction, *P*<5.0 × 10^−6^) eMAGMA gene-based results

**Table S4:** Significant (Bonferroni correction, *P*<1.7 × 10^−5^) S-PrediXcan gene-based results

**Table S5:** Overlap between significant MAGMA and eMAGMA gene-based results and the corresponding S-PrediXcan associations

**Table S6:** Overlap between significant eMAGMA and S-PrediXcan results

**Table S7:** MAGMA gene-set analyses before and after the removal of the MHC region

**Table S8:** eMAGMA gene-set analyses before and after the removal of the MHC region

**Table S9:** Biological pathway analysis of gene co-expression modules enriched with major depression genome-wide association signals

**Table S10:** Biological pathway analysis of all genes from modules enriched with MAGMA gene-based associations

**Figure S1:**
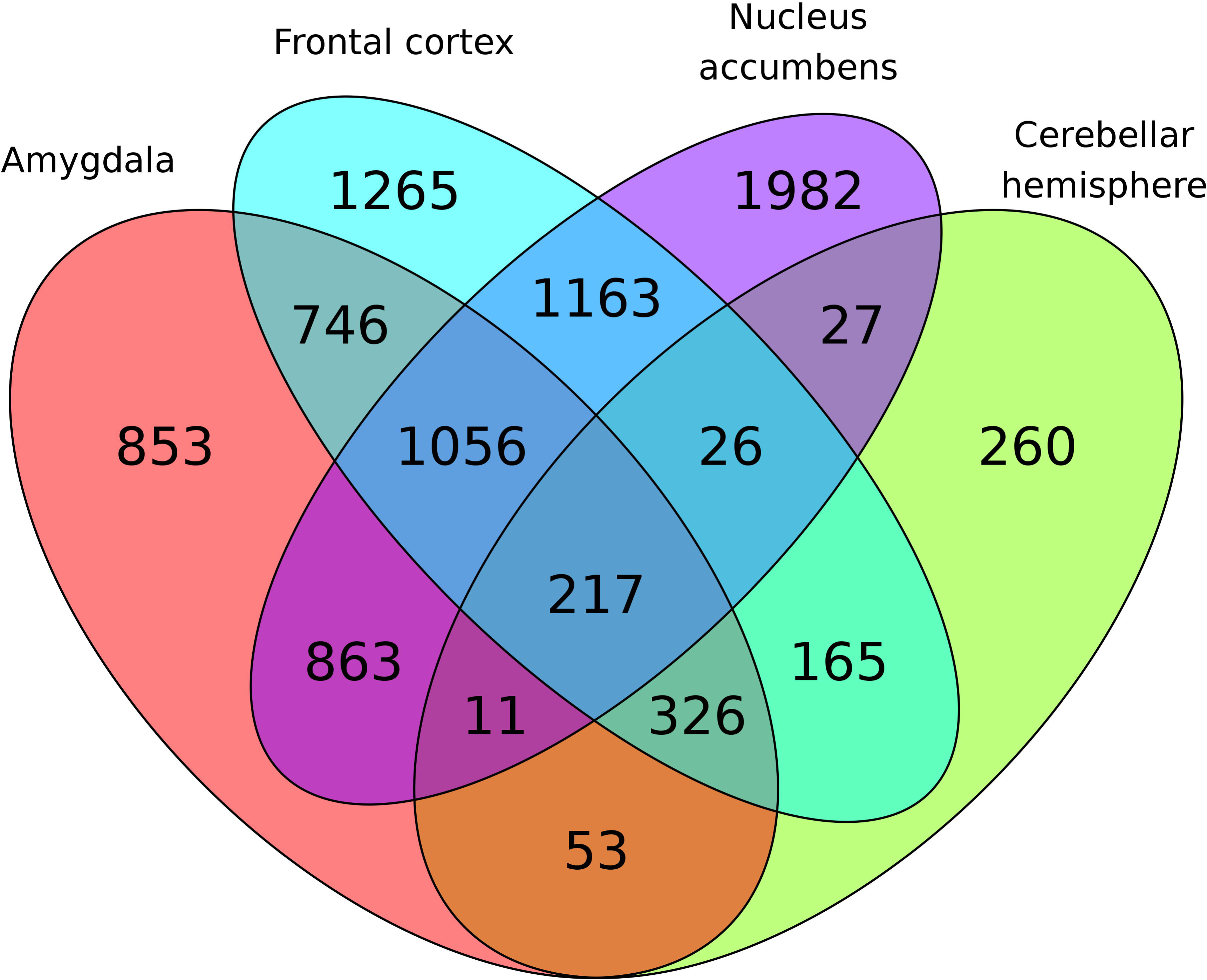
Overlap between genes within gene co-expression modules enriched with major depression signals from the MAGMA gene-based test

