## Supplementary Material 1 for "A Gene Co-expression Network-based Analysis of Multiple Brain Tissues Reveals Novel Genes and Molecular Pathways Underlying Major Depression"

Naomi R Wray* 1, 2

Stephan Ripke* 3, 4, 5

Manuel Mattheisen* 6, 7, 8, 9

Maciej Trzaskowski* 1

Enda M Byrne 1

Abdel Abdellaoui 10

Mark J Adams 11

Esben Agerbo 9, 12, 13

Tracy M Air 14

Till F M Andlauer 15, 16

Silviu-Alin Bacanu 17

Marie Bækvad-Hansen 9, 18

Aartjan T F Beekman 19

Tim B Bigdeli 17, 20

Elisabeth B Binder 15, 21

Douglas H R Blackwood 11

Julien Bryois 22

Henriette N Buttenschøn 8, 9, 23

Jonas Bybjerg-Grauholm 9, 18

Na Cai 24, 25

Enrique Castelao 26

Jane Hvarregaard Christensen 7, 8, 9

Toni-Kim Clarke 11

Jonathan R I Coleman 27

Lucía Colodro-Conde 28

Baptiste Couvy-Duchesne 2, 29

Nick Craddock 30

Gregory E Crawford 31, 32

Gail Davies 33

Ian J Deary 33

Franziska Degenhardt 34, 35

Eske M Derks 28

Nese Direk 36, 37

Conor V Dolan 10

Erin C Dunn 38, 39, 40

Thalia C Eley 27

Valentina Escott-Price 41

Farnush Farhadi Hassan Kiadeh 42

Hilary K Finucane 43, 44

Jerome C Foo 45

Andreas J Forstner 34, 35, 46, 47

Josef Frank 45

Héléna A Gaspar 27

Michael Gill 48

Fernando S Goes 49

Scott D Gordon 28

Jakob Grove 7, 8, 9, 50

Lynsey S Hall 11, 51

Christine Søholm Hansen 9, 18

Thomas F Hansen 52, 53, 54

Stefan Herms 34, 35, 47

Ian B Hickie 55

Per Hoffmann 34, 35, 47

Georg Homuth 56

Carsten Horn 57

Jouke-Jan Hottenga 10

David M Hougaard 9, 18

Marcus Ising 58

Rick Jansen 19

Ian Jones 59

Lisa A Jones 60

Eric Jorgenson 61

James A Knowles 62

Isaac S Kohane 63, 64, 65

Julia Kraft 4

Warren W. Kretzschmar 66

Jesper Krogh 67

Zoltán Kutalik 68, 69

Yihan Li 66

Penelope A Lind 28

Donald J MacIntyre 70, 71

Dean F MacKinnon 49

Robert M Maier 2

Wolfgang Maier 72

Jonathan Marchini 73

Hamdi Mbarek 10

Patrick McGrath 74

Peter McGuffin 27

Sarah E Medland 28

Divya Mehta 2, 75

Christel M Middeldorp 10, 76, 77

Evelin Mihailov 78

Yuri Milaneschi 19

Lili Milani 78

Francis M Mondimore 49

Grant W Montgomery 1

Sara Mostafavi 79, 80

Niamh Mullins 27

Matthias Nauck 81, 82

Bernard Ng 80

Michel G Nivard 10

Dale R Nyholt 83

Paul F O'Reilly 27

Hogni Oskarsson 84

Michael J Owen 59

Jodie N Painter 28

Carsten Bøcker Pedersen 9, 12, 13

Marianne Giørtz Pedersen 9, 12, 13

Roseann E. Peterson 17, 85

Erik Pettersson 22

Wouter J Peyrot 19

Giorgio Pistis 26

Danielle Posthuma 86, 87

Jorge A Quiroz 88

Per Qvist 7, 8, 9

John P Rice 89

Brien P. Riley 17

Margarita Rivera 27, 90

Saira Saeed Mirza 36

Robert Schoevers 91

Eva C Schulte 92, 93

Ling Shen 61

Jianxin Shi 94

Stanley I Shyn 95

Engilbert Sigurdsson 96

Grant C B Sinnamon 97

Johannes H Smit 19

Daniel J Smith 98

Hreinn Stefansson 99

Stacy Steinberg 99

Fabian Streit 45

Jana Strohmaier 45

Katherine E Tansey 100

Henning Teismann 101

Alexander Teumer 102

Wesley Thompson 9, 53, 103, 104

Pippa A Thomson 105

Thorgeir E Thorgeirsson 99

Matthew Traylor 106

Jens Treutlein 45

Vassily Trubetskoy 4

André G Uitterlinden 107

Daniel Umbricht 108

Sandra Van der Auwera 109

Albert M van Hemert 110

Alexander Viktorin 22

Peter M Visscher 1, 2

Yunpeng Wang 9, 53, 104

Bradley T. Webb 111

Shantel Marie Weinsheimer 9, 53

Jürgen Wellmann 101

Gonneke Willemsen 10

Stephanie H Witt 45

Yang Wu 1

Hualin S Xi 112

Jian Yang 2, 113

Futao Zhang 1

Volker Arolt 114

Bernhard T Baune 115

Klaus Berger 101

Dorret I Boomsma 10

Sven Cichon 34, 47, 116, 117

Udo Dannlowski 114

EJC de Geus 10, 118

J Raymond DePaulo 49

Enrico Domenici 119

Katharina Domschke 120

Tõnu Esko 5, 78

Hans J Grabe 109

Steven P Hamilton 121

Caroline Hayward 122

Andrew C Heath 89

Kenneth S Kendler 17

Stefan Kloiber 58, 123, 124

Glyn Lewis 125

Qingqin S Li 126

Susanne Lucae 58

Pamela AF Madden 89

Patrik K Magnusson 22

Nicholas G Martin 28

Andrew M McIntosh 11, 33

Andres Metspalu 78, 127

Ole Mors 9, 128

Preben Bo Mortensen 8, 9, 12, 13

Bertram Müller-Myhsok 15, 16, 129

Merete Nordentoft 9, 130

Markus M Nöthen 34, 35

Michael C O'Donovan 59

Sara A Paciga 131

Nancy L Pedersen 22

Brenda WJH Penninx 19

Roy H Perlis 38, 132

David J Porteous 105

James B Potash 133

Martin Preisig 26

Marcella Rietschel 45

Catherine Schaefer 61

Thomas G Schulze 45, 93, 134, 135, 136

Jordan W Smoller 38, 39, 40

Kari Stefansson 99, 137

Henning Tiemeier 36, 138, 139

Rudolf Uher 140

Henry Völzke 102

Myrna M Weissman 74, 141

Thomas Werge 9, 53, 142

Cathryn M Lewis 27, 143

Douglas F Levinson 144

Gerome Breen 27, 145

Anders D Børglum 7, 8, 9

Patrick F Sullivan 22, 146, 147,

1, Institute for Molecular Bioscience, The University of Queensland, Brisbane, QLD, AU

2, Queensland Brain Institute, The University of Queensland, Brisbane, QLD, AU

3, Analytic and Translational Genetics Unit, Massachusetts General Hospital, Boston, MA, US

4, Department of Psychiatry and Psychotherapy, Universitätsmedizin Berlin Campus Charité Mitte, Berlin, DE

5, Medical and Population Genetics, Broad Institute, Cambridge, MA, US

6, Centre for Psychiatry Research, Department of Clinical Neuroscience, Karolinska Institutet, Stockholm, SE

7, Department of Biomedicine, Aarhus University, Aarhus, DK

8, iSEQ, Centre for Integrative Sequencing, Aarhus University, Aarhus, DK

9, iPSYCH, The Lundbeck Foundation Initiative for Integrative Psychiatric Research,, DK

10, Dept of Biological Psychology & EMGO+ Institute for Health and Care Research, Vrije Universiteit Amsterdam, Amsterdam, NL

11, Division of Psychiatry, University of Edinburgh, Edinburgh, GB

12, Centre for Integrated Register-based Research, Aarhus University, Aarhus, DK

13, National Centre for Register-Based Research, Aarhus University, Aarhus, DK

14, Discipline of Psychiatry, University of Adelaide, Adelaide, SA, AU

15, Department of Translational Research in Psychiatry, Max Planck Institute of Psychiatry, Munich, DE

16, Munich Cluster for Systems Neurology (SyNergy), Munich, DE

17, Department of Psychiatry, Virginia Commonwealth University, Richmond, VA, US

18, Center for Neonatal Screening, Department for Congenital Disorders, Statens Serum Institut, Copenhagen, DK

19, Department of Psychiatry, Vrije Universiteit Medical Center and GGZ inGeest, Amsterdam, NL

20, Virginia Institute for Psychiatric and Behavior Genetics, Richmond, VA, US

21, Department of Psychiatry and Behavioral Sciences, Emory University School of Medicine, Atlanta, GA, US

22, Department of Medical Epidemiology and Biostatistics, Karolinska Institutet, Stockholm, SE

23, Department of Clinical Medicine, Translational Neuropsychiatry Unit, Aarhus University, Aarhus, DK

24, Human Genetics, Wellcome Trust Sanger Institute, Cambridge, GB

25, Statistical genomics and systems genetics, European Bioinformatics Institute (EMBL-EBI), Cambridge, GB

26, Department of Psychiatry, University Hospital of Lausanne, Prilly, Vaud, CH

27, Social Genetic and Developmental Psychiatry Centre, King's College London, London, GB

28, Genetics and Computational Biology, QIMR Berghofer Medical Research Institute, Brisbane, QLD, AU

29, Centre for Advanced Imaging, The University of Queensland, Brisbane, QLD, AU

30, Psychological Medicine, Cardiff University, Cardiff, GB

31, Center for Genomic and Computational Biology, Duke University, Durham, NC, US

32, Department of Pediatrics, Division of Medical Genetics, Duke University, Durham, NC, US

33, Centre for Cognitive Ageing and Cognitive Epidemiology, University of Edinburgh, Edinburgh, GB

34, Institute of Human Genetics, University of Bonn, Bonn, DE

35, Life&Brain Center, Department of Genomics, University of Bonn, Bonn, DE

36, Epidemiology, Erasmus MC, Rotterdam, Zuid-Holland, NL

37, Psychiatry, Dokuz Eylul University School Of Medicine, Izmir, TR

38, Department of Psychiatry, Massachusetts General Hospital, Boston, MA, US

39, Psychiatric and Neurodevelopmental Genetics Unit (PNGU), Massachusetts General Hospital, Boston, MA, US

40, Stanley Center for Psychiatric Research, Broad Institute, Cambridge, MA, US

41, Neuroscience and Mental Health, Cardiff University, Cardiff, GB

42, Bioinformatics, University of British Columbia, Vancouver, BC, CA

43, Department of Epidemiology, Harvard T.H. Chan School of Public Health, Boston, MA, US

44, Department of Mathematics, Massachusetts Institute of Technology, Cambridge, MA, US

45, Department of Genetic Epidemiology in Psychiatry, Central Institute of Mental Health,  Medical Faculty Mannheim, Heidelberg University, Mannheim, Baden-Württemberg, DE

46, Department of Psychiatry (UPK), University of Basel, Basel, CH

47, Human Genomics Research Group, Department of Biomedicine, University of Basel, Basel, CH

48, Department of Psychiatry, Trinity College Dublin, Dublin, IE

49, Psychiatry & Behavioral Sciences, Johns Hopkins University, Baltimore, MD, US

50, Bioinformatics Research Centre, Aarhus University, Aarhus, DK

51, Institute of Genetic Medicine, Newcastle University, Newcastle upon Tyne, GB

52, Danish Headache Centre, Department of Neurology, Rigshospitalet, Glostrup, DK

53, Institute of Biological Psychiatry, Mental Health Center Sct. Hans, Mental Health Services Capital Region of Denmark, Copenhagen, DK

54, iPSYCH, The Lundbeck Foundation Initiative for Psychiatric Research, Copenhagen, DK

55, Brain and Mind Centre, University of Sydney, Sydney, NSW, AU

56, Interfaculty Institute for Genetics and Functional Genomics, Department of Functional Genomics, University Medicine and Ernst Moritz Arndt University Greifswald, Greifswald, Mecklenburg-Vorpommern, DE

57, Roche Pharmaceutical Research and Early Development, Pharmaceutical Sciences, Roche Innovation Center Basel, F. Hoffmann-La Roche Ltd, Basel, CH

58, Max Planck Institute of Psychiatry, Munich, DE

59, MRC Centre for Neuropsychiatric Genetics and Genomics, Cardiff University, Cardiff, GB

60, Department of Psychological Medicine, University of Worcester, Worcester, GB

61, Division of Research, Kaiser Permanente Northern California, Oakland, CA, US

62, Psychiatry & The Behavioral Sciences, University of Southern California, Los Angeles, CA, US

63, Department of Biomedical Informatics, Harvard Medical School, Boston, MA, US

64, Department of Medicine, Brigham and Women's Hospital, Boston, MA, US

65, Informatics Program, Boston Children's Hospital, Boston, MA, US

66, Wellcome Trust Centre for Human Genetics, University of Oxford, Oxford, GB

67, Department of Endocrinology at Herlev University Hospital, University of Copenhagen, Copenhagen, DK

68, Institute of Social and Preventive Medicine (IUMSP), University Hospital of Lausanne, Lausanne, VD, CH

69, Swiss Institute of Bioinformatics, Lausanne, VD, CH

70, Division of Psychiatry, Centre for Clinical Brain Sciences, University of Edinburgh, Edinburgh, GB

71, Mental Health, NHS 24, Glasgow, GB

72, Department of Psychiatry and Psychotherapy, University of Bonn, Bonn, DE

73, Statistics, University of Oxford, Oxford, GB

74, Psychiatry, Columbia University College of Physicians and Surgeons, New York, NY, US

75, School of Psychology and Counseling, Queensland University of Technology, Brisbane, QLD, AU

76, Child and Youth Mental Health Service, Children's Health Queensland Hospital and Health Service, South Brisbane, QLD, AU

77, Child Health Research Centre, University of Queensland, Brisbane, QLD, AU

78, Estonian Genome Center, University of Tartu, Tartu, EE

79, Medical Genetics, University of British Columbia, Vancouver, BC, CA

80, Statistics, University of British Columbia, Vancouver, BC, CA

81, DZHK (German Centre for Cardiovascular Research), Partner Site Greifswald, University Medicine, University Medicine Greifswald, Greifswald, Mecklenburg-Vorpommern, DE

82, Institute of Clinical Chemistry and Laboratory Medicine, University Medicine Greifswald, Greifswald, Mecklenburg-Vorpommern, DE

83, Institute of Health and Biomedical Innovation, Queensland University of Technology, Brisbane, QLD, AU

84, Humus, Reykjavik, IS

85, Virginia Institute for Psychiatric & Behavioral Genetics, Virginia Commonwealth University, Richmond, VA, US

86, Clinical Genetics, Vrije Universiteit Medical Center, Amsterdam, NL

87, Complex Trait Genetics, Vrije Universiteit Amsterdam, Amsterdam, NL

88, Solid Biosciences, Boston, MA, US

89, Department of Psychiatry, Washington University in Saint Louis School of Medicine, Saint Louis, MO, US

90, Department of Biochemistry and Molecular Biology II, Institute of Neurosciences, Center for Biomedical Research, University of Granada, Granada, ES

91, Department of Psychiatry, University of Groningen, University Medical Center Groningen, Groningen, NL

92, Department of Psychiatry and Psychotherapy, Medical Center of the University of Munich, Campus Innenstadt, Munich, DE

93, Institute of Psychiatric Phenomics and Genomics (IPPG), Medical Center of the University of Munich, Campus Innenstadt, Munich, DE

94, Division of Cancer Epidemiology and Genetics, National Cancer Institute, Bethesda, MD, US

95, Behavioral Health Services, Kaiser Permanente Washington, Seattle, WA, US

96, Faculty of Medicine, Department of Psychiatry, University of Iceland, Reykjavik, IS

97, School of Medicine and Dentistry, James Cook University, Townsville, QLD, AU

98, Institute of Health and Wellbeing, University of Glasgow, Glasgow, GB

99, deCODE Genetics / Amgen, Reykjavik, IS

100, College of Biomedical and Life Sciences, Cardiff University, Cardiff, GB

101, Institute of Epidemiology and Social Medicine, University of Münster, Münster, Nordrhein-Westfalen, DE

102, Institute for Community Medicine, University Medicine Greifswald, Greifswald, Mecklenburg-Vorpommern, DE

103, Department of Psychiatry, University of California, San Diego, San Diego, CA, US

104, KG Jebsen Centre for Psychosis Research, Norway Division of Mental Health and Addiction, Oslo University Hospital, Oslo, NO

105, Medical Genetics Section, CGEM, IGMM, University of Edinburgh, Edinburgh, GB

106, Clinical Neurosciences, University of Cambridge, Cambridge, GB

107, Internal Medicine, Erasmus MC, Rotterdam, Zuid-Holland, NL

108, Roche Pharmaceutical Research and Early Development, Neuroscience, Ophthalmology and Rare Diseases Discovery & Translational Medicine Area, Roche Innovation Center Basel, F. Hoffmann-La Roche Ltd, Basel, CH

109, Department of Psychiatry and Psychotherapy, University Medicine Greifswald, Greifswald, Mecklenburg-Vorpommern, DE

110, Department of Psychiatry, Leiden University Medical Center, Leiden, NL

111, Virginia Institute for Psychiatric & Behavioral Genetics, Virginia Commonwealth University, Richmond, VA, US

112, Computational Sciences Center of Emphasis, Pfizer Global Research and Development, Cambridge, MA, US

113, Institute for Molecular Bioscience; Queensland Brain Institute, The University of Queensland, Brisbane, QLD, AU

114, Department of Psychiatry, University of Münster, Münster, Nordrhein-Westfalen, DE

115, Department of Psychiatry, Melbourne Medical School, University of Melbourne, Melbourne, AU

116, Institute of Medical Genetics and Pathology, University Hospital Basel, University of Basel, Basel, CH

117, Institute of Neuroscience and Medicine (INM-1), Research Center Juelich, Juelich, DE

118, Amsterdam Public Health Institute, Vrije Universiteit Medical Center, Amsterdam, NL

119, Centre for Integrative Biology, Università degli Studi di Trento, Trento, Trentino-Alto Adige, IT

120, Department of Psychiatry and Psychotherapy, Medical Center, University of Freiburg, Faculty of Medicine, University of Freiburg, Freiburg, DE

121, Psychiatry, Kaiser Permanente Northern California, San Francisco, CA, US

122, Medical Research Council Human Genetics Unit, Institute of Genetics and Molecular Medicine, University of Edinburgh, Edinburgh, GB

123, Department of Psychiatry, University of Toronto, Toronto, ON, CA

124, Centre for Addiction and Mental Health, Toronto, ON, CA

125, Division of Psychiatry, University College London, London, GB

126, Neuroscience Therapeutic Area, Janssen Research and Development, LLC, Titusville, NJ, US

127, Institute of Molecular and Cell Biology, University of Tartu, Tartu, EE

128, Psychosis Research Unit, Aarhus University Hospital, Risskov, Aarhus, DK

129, University of Liverpool, Liverpool, GB

130, Mental Health Center Copenhagen, Copenhagen Universtity Hospital, Copenhagen, DK

131, Human Genetics and Computational Biomedicine, Pfizer Global Research and Development, Groton, CT, US

132, Psychiatry, Harvard Medical School, Boston, MA, US

133, Psychiatry, University of Iowa, Iowa City, IA, US

134, Department of Psychiatry and Behavioral Sciences, Johns Hopkins University, Baltimore, MD, US

135, Department of Psychiatry and Psychotherapy, University Medical Center Göttingen, Goettingen, Niedersachsen, DE

136, Human Genetics Branch, NIMH Division of Intramural Research Programs, Bethesda, MD, US

137, Faculty of Medicine, University of Iceland, Reykjavik, IS

138, Child and Adolescent Psychiatry, Erasmus MC, Rotterdam, Zuid-Holland, NL

139, Psychiatry, Erasmus MC, Rotterdam, Zuid-Holland, NL

140, Psychiatry, Dalhousie University, Halifax, NS, CA

141, Division of Epidemiology, New York State Psychiatric Institute, New York, NY, US

142, Department of Clinical Medicine, University of Copenhagen, Copenhagen, DK

143, Department of Medical & Molecular Genetics, King's College London, London, GB

144, Psychiatry & Behavioral Sciences, Stanford University, Stanford, CA, US

145, NIHR Maudsley Biomedical Research Centre, King's College London, London, GB

146, Genetics, University of North Carolina at Chapel Hill, Chapel Hill, NC, US

147, Psychiatry, University of North Carolina at Chapel Hill, Chapel Hill, NC, US

Version 5. 2018-07-17
